# Ribbit: Accurate identification and annotation of complex tandem repeat sequences in genomes

**DOI:** 10.1101/2025.02.06.636828

**Authors:** Akshay Kumar Avvaru, Anukrati Sharma, Divya Tej Sowpati

## Abstract

DNA tandem repeats (TRs) are crucial for genomic functions like protein binding, chromatin modulation, splicing, and gene regulation. Abnormal length variations in TRs, especially expansions, are associated with over 60 neurodegenerative diseases. The function and stability of a TR locus is dependent on its sequencing composition and purity. Recent studies report the disease-causing propensity of non-canonical motif expansions in TR loci, and highlight the intricate polymorphism dynamics in complex loci encompassing adjacent, overlapping, and nested TRs. These reports emphasize the need for precise definition and motif decomposition of TR loci. To address this, we present Ribbit, a tool that accurately and efficiently identifies and annotates TR loci in a genome. Ribbit uses 2-bit representation of DNA sequences for rapid identification of TRs of 2–100 bp motif size and resolves complex TR structures. Ribbit efficiently handles imperfections such as indels and substitutions, providing insights into nested and compound TR relationships through detailed motif decomposition. Comparative analyses using simulated data show Ribbit outperforms existing tools like Dot2dot and TRF in terms of runtime and accuracy. Ribbit reports TR loci in the human genome with lower redundancy than TRF and provides resolved TR regions comparable to variation clusters reported in recent catalogues. Therefore, Ribbit can be leveraged to understand the evolution and biology of complex TR regions in large genomes.

## Introduction

Tandem repeats are genomic sequences with contiguous copies of a DNA motif. TRs are classified based on the size of the motif as short tandem repeats (STRs) with 2-6 bps long motifs and variable number tandem repeats (VNTRs) with motifs of 7 bps and above. Tandem repeats show a characteristic length polymorphism with changes in the number of unit repetitions of a locus. These are highly polymorphic in nature with mutation rates in the range of 10^−6^ to 10^−4^ per generation (Chiu et al., 2024a). Abnormal variations in length, particularly expansions, of certain TR loci are causatively associated with >60 neurodegenerative diseases in humans including Fragile X syndrome, Huntington’s disease, and multiple ataxias. Many TR loci are shown to have functional roles including protein binding, modulating chromatin landscape, alter splicing and regulate gene expression. Profiling variation at TR loci has been strategically difficult due to ambiguous sequence mapping and alignment besides limitations of sequencing technologies. Innovations mitigating sequencing errors, improving read lengths and robust computational methods improved accuracy of alignment, assembly and genotyping at repetitive regions. Studying variation at tandem repeat loci in large population cohorts such as the UK BioBank has become common, on par with SNVs and SVs. Large scale population studies expanded the catalogue of polymorphic TR loci and revealed loci associated with quantitative traits including transcriptomic and methylation profiles. Long read genome sequencing aided the discovery of novel disease associated TR loci including many VNTRs and STRs with unconventional mutation mechanisms. Besides expansions and contractions of canonical motifs in a TR, inclusion of non-canonical motifs and their expansions are discovered to be pathogenic. Example, the presence of non-canonical motifs AGGGC, AAGGC and AGAGG in the *RFC1* tandem repeat locus is pathogenic while expansions of the canonical motifs AAAAG and AAAGG are benign (Natalia Dominik et al., 2023 Brain; Satoko Miyatake et al., 2022 Brain). These unconventional variations in tandem repeats and association to disease and function draw importance to understanding the sequence complexity of a TR locus.

Ambiguity in TR definitions across different studies made studies inconsistent and also unreproducible, raising the need for a standard catalog of TR loci. Multiple groups have proposed an ideal set of TR loci in the human genome to be used across population studies (Adam C. English et al. 2024, Readman Chiu et al. 2024, Ben Weisburd et al. 2024). Individually these studies have identified the importance of categorizing the loci into simple or complex tandem repeats. A complex TR locus constitutes multiple adjacent or overlapping TRs, making it difficult to identify the locus boundaries and therefore leads to ambiguity in variant detection. Repeats cataloged under GIAB benchmark are classified as overlapping, isolated, nested and adjacent where 68.9% of repeats had non overlapping annotations, 19% had parent/nested annotations and the rest as overlapping [Adam English 2023]. Weisburd et al. have proposed a data driven method to define complex TR loci and termed them variation clusters. Tandem repeat loci with higher frequency of observed polymorphisms, from population studies, in their neighborhood are extended often to include neighboring tandem repeats forming a variation cluster (VC). The authors identified 273,112 VCs that cumulatively overlapped 744,458 TRs, from a catalog of 4,863,043 TRs. While building a comprehensive catalog of tandem repeats with a total of 18,759,399 loci Readman Chiu et al. have only retained TR locus with higher alignment score between a pair of overlapping loci. These catalogs are essentially built by merging existing sets of TR loci either identified to be polymorphic in population studies or are generated based on defined parameters using TR identification tools. These studies revealed complex TR loci that constitute TRs with nested-parent relationships, overlapping TRs or single locus with dual interpretations of periodicities and consensus motif. TR identification tools have come a long way in identifying and annotating simple TR loci delineating coordinate boundaries, calculating periodicity, resolving longer motifs and determining consensus motif. But, methods have not been developed or extended for identification and annotation of complex tandem repeat loci. Tools unable to resolve and report complex TR regions including nested, overlapping or multiple arrays of one motif, gets difficult to genotype such regions as most genotyping tools work best with pure or nearly pure repeat loci [Ben Weisburd 2024].

Ambiguity in tandem repeat (TR) definitions across studies has led to inconsistencies and irreproducibility, highlighting the need for a standardized catalog of TR loci. Many studies aim to categorize TR loci as either simple or complex, with complex loci comprising multiple adjacent or overlapping repeats. These complexities make it challenging to define locus boundaries, introducing ambiguity in variant detection (Adam C. English et al. 2024, Readman Chiu et al. 2024, Ben Weisburd et al. 2024). Weisburd et al. proposed a data-driven approach to address this issue by defining complex TR loci as “variation clusters, which are highly polymorphic. TR identification tools have come a long way in identifying and annotating simple TR loci delineating coordinate boundaries, calculating periodicity, resolving longer motifs and determining consensus motif. But, methods have not been developed or extended for identification and annotation of complex tandem repeat loci. Unresolved TR regions with nested, overlapping or multiple arrays of one motif, are difficult to genotype as most genotyping tools work best with pure or nearly pure repeat locations. This challenge is universal across genomes as complex TRs can provide better insights about the repeat profile of an organism.

We here propose Ribbit, developed specifically to resolve complex tandem repeats. Ribbit processes a sequence for potential tandem repeat of all periodicities in three stages, relaxing the threshold purity at each stage. In the first stage Ribbit looks for perfect TR sequences, the second stage allows mismatches between the motifs of a TR sequence and the third allows for indels between the motifs. The hierarchical identification allows for comparison of overlapping interpretations of a locus as a potential TR of different periodicities and retains all interpretations based on higher repeat purity. The potential TRs are identified from 2-bit representations of DNA sequences which completes this three stage identification of a sequence in comparable if not more efficient time and memory required than existing tools. We show the accuracy of Ribbit matches that of widely used TR identification tools, including TRF and Dot2dot. We present two case studies of complex TR loci where Ribbit resolves the loci retaining all valid TRs of higher purity within the locus. To tackle the reporting of complex TR loci we generate the outputs in two formats, 1) BED file having all the TRs as individual loci 2) TR catalog as a GTF file with relationships defined between TR loci which are part of a complex repeat.

## Methods

### Ribbit algorithm Overview

A dot-plot built from self-aligning a sequence containing a TR with periodicity *d* and *r* units results in a distinct pattern of matches along all the diagonals at a distance *du* (u □ {1, 2, …, r}) parallel to the central diagonal (Fig 1A). Ideally, identification of TR loci of all periodicities could be achieved by searching for stretches with matches along all the diagonals from a dot-plot. Ribbit is based on this particular idea of identifying TR sequences from dot-plot diagonals. A diagonal at distance *d* from the central diagonal is analogous to an alignment of a sequence *S*, of length *s*, with the sequence generated by shifting *S* by *d* nucleotides. Ribbit uses 2-bit representation of DNA sequences to generate these diagonals using two basic bit operations 1) *bitshift*: generates *S*_*d*_ i.e., *S* shifted by *d* nucleotides 2) *XOR* - compares *S* with *S*_*d*_. The comparison yields a bit-sequence, *SXOR*_*d*_, of the length *s* with 1s denoting matches and 0s denoting the mismatches along the diagonal *d*. Continuous matches along *SXOR*_*d*_ denote the presence of a TR of periodicity *d*. Mismatches between motifs of a TR interrupt the matches *SXOR*_*d*_. Insertions and deletions of size δ between the motifs show a stretch of 0s along *SXOR*_*d*_ but will be complemented with 1s in the *u+*δ and *u-*δ shift diagonals respectively: An insertion of size 1 results in a maximum of *d+1* 0s in the *SXOR*_*d*_ but will have a continuous set of 1s at the same position in *SXOR*_*d+1*_. Similar.y, a deletion of 1 base will show up as a *d-1* 0s in *SXOR*_*d*_ but will have *u-1* 1s in the same positions in *SXOR*_*d-1*_. Ribbit leverages these rules to identify potential TRs from a sequence.

**Figure 1.**
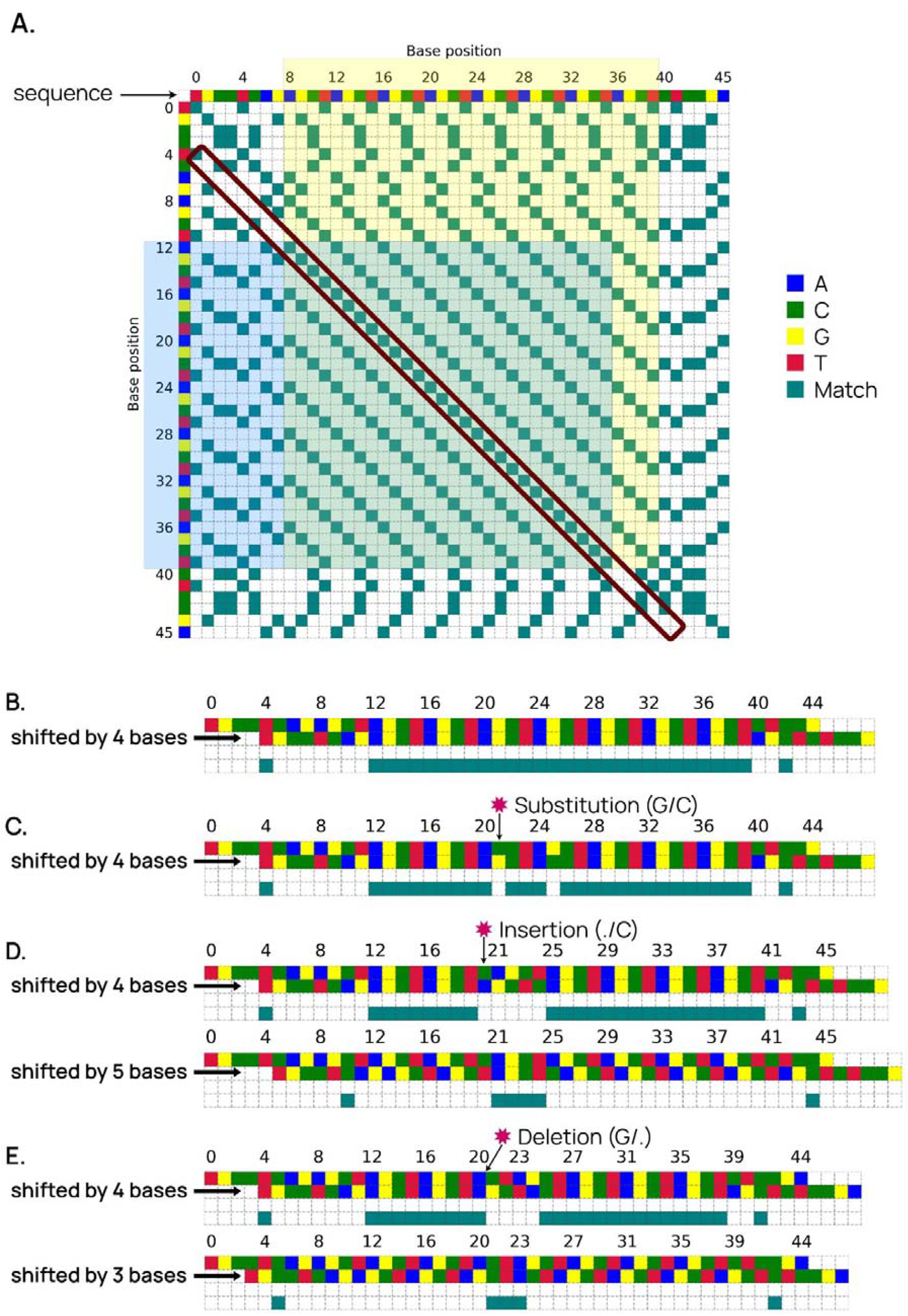
Core algorithm of Ribbit explained. A) The dotplot of a 45bp DNA sequence with a TR of motif AGCT repeated 8 times from position 8 to 39. Dot plot indicates matches along the 4th diagonal exactly at the repeat coordinates. B) Depiction of 4 bases shifted self-alignment of the sequence indicating matches identical to the 4th diagonal in the dot plot. C) Shows the interruption in matches with a mismatch present at position 21 in the sequence. D) Shows the interruption in matches along the 4th shift self-alignment due to an insertion at position 20. These mismatches are observed to be compensated along the 5th shift self-alignment. E) Shows the interruption in matches along the 4th shift self-alignment due to an deletion at position 21. These mismatches are observed to be compensated along the 3rd shift self-alignment.

**Figure 2.**
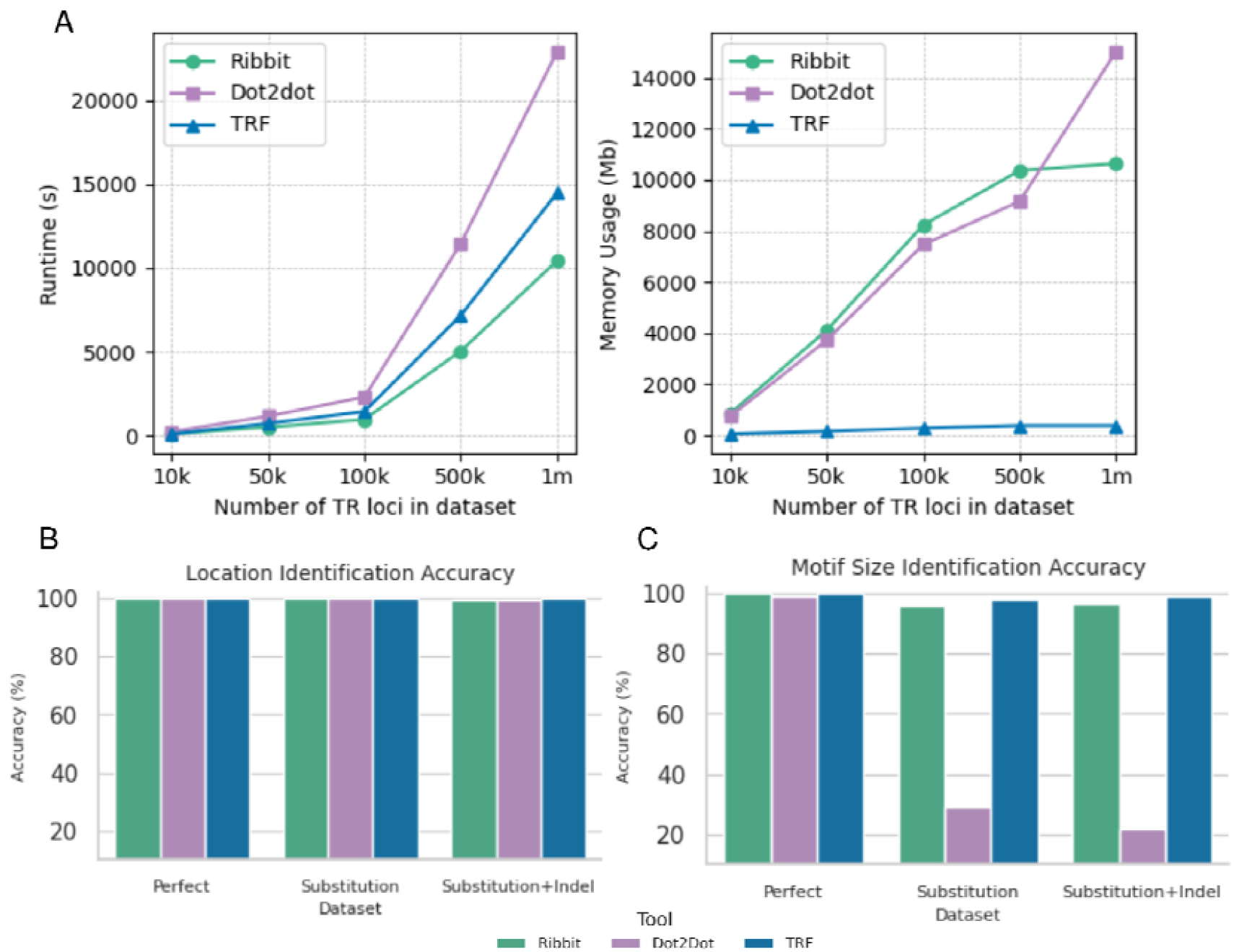
Performance of tools on Simulated datasets with Substitution and Indels across all sample sizes. **A)** Runtime comparison and Memory Usage comparison. **B,C)** Comparison of location identification and motif size identification accuracy across different purity levels in simulated datasets with a sample size of 1 million.

### 2-bit conversion of the DNA sequence

Ribbit converts DNA sequence into its binary representation to minimize the computational overhead associated with string operations. The dynamic bitset data structure from the boost library is used to store the converted 2-bit DNA sequences. This bitset data structure dynamically allocates memory for the bitset at runtime eluding the restrictions that are imposed at compile time. The nucleotide sequence is converted into 2-bit representation using a lexicographic order of nucleotide to bit mapping (A=00, C=01, G=10, T=11). The sequence is represented as a combination of two bitsets, one holding the left bit of each nucleotide and other holding the right (Fig 1A). A third bitset of the same length is used to mark all the Ns in the sequence. Ribbit also builds a dot matrix representation of the sequence by adapting the data structure developed in Dot2dot. Individual bitsets are built to track the positions of each nucleotide (A, C, G, T). To represent the complete sequence as a dot matrix a vector of pointers is built, with each pointer pointing to the bitset corresponding to the nucleotide at the position (Fig.1B).

### Using bitsets to identify tandem repeat sequences

The core algorithm of Ribbit is better understood through the dot plot representation used for visualizing repeats in a DNA sequence (Figure 1A). In a dot plot built from a self alignment of a sequence, the diagonals correspond to alignment of the sequence with a frame shifted version of itself (Figure 1B). Matches along the diagonal at the parallel distance of *d* from the central diagonal indicate the presence of a TR of periodicity *d*. To build these diagonals Ribbit uses the 2-bit representation of DNA. The bit sequence data structure allows for the usage of computationally efficient bit shift and XOR operations. Ribbit builds these diagonals by converting DNA sequences to 2-bit format followed by bit shift and XOR operations. Shifting the bit sequence by *d* bits and performing an XOR with the original bit sequence, aligns every nucleotide at position *i* to a nucleotide at position *i* + *d*. XOR operation indicates matches by 0, hence the resultant bit sequence is flipped to indicate matches by 1.

### Interpreting motif mismatches and indels along dot-plot diagonals

A TR of periodicity *d* is indicated by matches along the diagonal of distance *d* from the central diagonal in the dot-plot. Substitutions or mismatches between adjacent motifs are indicated by a 0s along the *d* diagonal (Figure 1C). Indels between motifs disrupt the frame of the tandem repeat. While they are also indicated by 0s along the diagonal they are complemented by matches along the diagonal ± □ distance from the TR diagonal, where □ denotes the length of the indel (Figure 1D, 1E). Ribbit uses this logic and complements shift XOR of *d* with shift XORs of *d* ± □.

### Using anchor bitsets to account for indels

As explained in Figure 1C, indels of sizes □ between motifs shows up as matches in *d* ± □ shift XORs. To account for indels we complement the *d* shift XOR with small continuous matches from *d* - □ to *d* + □ shift XORs. For this we build the anchor shift XORs. Anchor shift XOR of *d* shift is the shift XOR bitset with only continuous matches of at least □ and maximum *d* retained. Matches of length beyond this range are set to 0. The maximum threshold of *d* continuous matches filters the native repeats with periodicity *d*. These smaller stretches of matches from the *d* shift XOR could now be used to evaluate insertions in TRs of *d* - 1 periodicity and deletions in TRs of *d* + 1 periodicity.

### Stratified identification of TR seed locus

One of the main problems of imperfect TR identification is redundant reinterpretations of the same locus. One reason is degenerate interpretation of periodicity; a TR with periodicity *d* shows matches along the diagonal corresponding to *d, 2d, 3d* and further integer multiples of *d*. The same locus is hence reported as TR with periodicities for all integer multiples of *d*. Another reason for redundancy is multiple periodicity annotations due to allowed mismatches and indels between the motifs. Ribbit tackles these issues using a ranked evaluation of the TR locus by stepwise inclusion of mismatches and indels. Ribbit scans the shift XORs of all desired motif sizes with window size *w* with complete matches to identify stretches of perfect, uninterrupted TR sequences. Loci identified in the *d* shift XOR are stored as a seed TR locus of periodicity *d*. Seeds annotated with periodicity *d* are omitted if an overlapping locus is annotated as a seed with periodicity that is a factor of *d*. This first scan of shift XORs for all desired periodicities results in identification of perfect TR stretches.

The shift XORs are next scanned again with a window size of *w* for windows with at least *w* - 1 matches, which picks up TR loci with mismatches between motifs. The seeds picked up in this step are compared with the perfect TR seeds and merged where periodicity annotations match. The seeds picked up with mismatches allowed are always larger than the perfect stretches hence either extend the range of the perfect seed or merge multiple seeds. Ribbit compares the seeds picked up from the first and second steps strategically to achieve proper extension and merging (elaborated in supplementary note).

To further include indels Ribbit complements the *d* shift XOR with anchor shift XORs of *d* - □ to *d* + □ shift XORs. The new anchored shift XORs are created with OR operations between the *d* shift XOR and anchor shift XORs of *d* - □ to *d* + □ shift XORs. The new anchored shift XOR of *d* shift is now scanned with a window size of *w* and threshold matches of *w* - 1. Ribbit now picks up TR seeds of periodicity *d* with included mismatches and indels allowed between the motifs.

### Ranked evaluation of TR locus seeds

The seed identification takes place in three stages and each stage deals with a different level of purity of the repetitive location. Ribbit begins with scanning for perfect TR loci which are ranked as the highest priority seeds because they are least likely to be redundant, and their periodicity can be interpreted confidently. The seeds picked from the first stage are stored within a vector with start, end coordinates, periodicity, and are given the rank *A*. At this stage redundant seeds are filtered to only select for seeds with atomic motif sizes. Ribbit then identifies seeds with mismatches, which are given the rank B. The seeds identified at this stage are dynamically compared with the seeds identified in the previous stage. Overlapping seeds with matching or periodicities with factor-multiple relation across both ranks are merged, with atomic periodicity retained. Overlapping TR seeds with periodicities *ds* and *dl*, where *ds* < *dl* and *dl* □ *ds*, are merged and annotated with periodicity *ds*, depending on the overlapping length and size ranges of *ds* and *dl*. This rule of shorter periodicity getting a priority when two seeds are merged is applied for both partial and complete overlaps. In case of overlapping seeds with nested-parent relationships, the nested seed is dropped if it has a lower rank. The rank helps Ribbit in choosing the best possible representation for the TR location in case of overlaps to reduce redundancy. Next, Ribbit identified TRs with indels, which are assigned the rank C. Similar to before, seeds identified at this stage are compared dynamically with previously identified seeds. To pick the most appropriate seed representation for a locus with overlapping TR interpretations across different ranks, ribbit carries out filtering or merging based on periodicities, ranks and length of overlap to get rid of redundant false representations of the locus. The core logic of filtering and merging strategy is that between the overlapping seeds the higher rank seed is given priority for deciding the periodicity of the TR locus and lower ranked seeds with different periodicity interpretations are dropped.

### Consensus motif identification

Seed loci are potential TR sequences designated based on high frequency nucleotide matches with a particular periodicity *d*. Each seed is now processed to define its sequence properties including the consensus motif sequence of the repeated units and the mismatches/indels between the unit motifs. Ribbit employs two different approaches for different classes of TRs based on the motif size. For seeds with periodicity *d* ≤ 10 referred to as microsatellites Ribbit uses KMP (knuth morris pratt) algorithm with a sliding window approach with window size *d*. In the KMP algorithm a motif sequence is tracked based on all the indices it is occurring in the seed sequence. Ribbit groups the indices for cyclical motifs under one representative motif. Motif groups with index ranges that span a user-defined length threshold without a gap between the indices larger than 3 motif lengths are designated as the final TR sequence of the representative motif. This approach makes sure that every coordinate span picked for a motif starts and ends with the complete motif. Also, this approach delineates coordinate ranges for TRs with identical periodicity but differing representative motifs, that are either overlapping or separated by short distance and merged together in a single seed. For TRs with periodicities *d* > 10 ribbit employs a dot matrix based approach to identify a consensus motif sequence. It examines stacked diagonals occurring at intervals equal to the motif size, specifically at seed indices. First, the dot matrix *D* is built for the seed sequence of length *s*. Ribbit then picks each k-mer of between indices *i* to *i* + *d* along the seed sequence where 0 ≤ *i ≤ s* - *d*. The similarity of this motif with the contiguous motif downstream is calculated by counting matches along the diagonal *D*[*i, i*+*d*] to *D*[*i*+*d, i*+2*d*]. Similarity with the contiguous motif upstream is calculated by counting matches along the diagonal *D*[*i, i*-*d*] to *D*[*i, i*+*d*]. Similarly, the similarity with all the contiguous motifs is calculated by counting matches along all corresponding diagonals. The motif with the highest calculated matches is considered as the consensus sequence.

The seed sequences are the potential location for TRs with certain imperfections. The algorithm utilizes known parameters to retrieve the DNA sequence corresponding to the seed bitset. The motif size of a repeating unit within a tandem repeat (TR) can be determined by observing the number of shifts required to achieve matches in the string. Ribbit processes seeds with shorter motif length (2-6) and longer motif lengths (beyond 6) with different algorithms. In the DNA sequence corresponding to the seed bitset the algorithm looks for the most frequent motif of the known motif size. Ribbit uses KMP (knuth morris pratt) algorithm with a sliding window approach for shorter motifs. Here, the window size is set equal to the motif size. The algorithm systematically analyzes the sequences, determining the frequency of each motif through keeping a count of occurrences of the windows. The most frequently occurring window is selected as the primary motif. In cases where two or more windows exhibit the same frequency and represent cyclic variations, the repeat class is designated as the motif. However, if the motifs are not cyclic variations but share equal frequencies, both motifs are retained for the formation of pseudo-perfect sequences, representing compound repeats. [about the remaining part of the seed where there is a recheck for another motif to look at compound repeats.

Ribbit uses a dot matrix approach to identify most frequent motifs for repeats with motif size longer than 6 base pairs. It searches for repeating patterns by examining the stacked diagonals occurring at intervals equal to the motif size. Ribbit computes the average count of matches for diagonals and takes the row interval with the maximum average count. The sequence within this row interval is taken as the motif for the TR.

### Measuring the purity

Post the identification of a consensus motif for a seed TR locus Ribbit evaluates the sequence for purity by comparing it with the ideal counterpart i.e., perfect repeat sequence. In this process, a pseudo-perfect TR, with a length equal to the seed sequence, is generated by repeating the consensus motif. This pseudo-perfect TR serves as a reference for evaluating the impurities within the original seed sequence. Ribbit employs the SSW C/C++ library, a Single Instruction Multiple Data (SIMD) implementation of the Smith-Waterman alignment, to efficiently perform these alignments(Zhao et al., 2013). The alignment of the seed sequence, extracted from the reference genome, and the pseudo perfect repeat yields reference/query coordinates, SW score and a CIGAR which are used to evaluate the impurities in the repeat. The purity of the TR locus is calculated as the number of matches divided by the total alignment length, which includes matches, mismatches & indel lengths.

### Generating Simulated Data

To generate the simulated data, we utilized a custom Python script specifically designed to model repeat loci with motif sizes ranging from 2 to 100 bases. To achieve a comprehensive representation of tandem repeats, we incorporated a randomized motif selection process from a list of all the possible combinations of nucleotides for a motif size. This ensured that the simulation captured the diversity of repeat structures observed in genomic sequences. To reflect realistic distributions of motif sizes, we first analyzed the human genome (hg38) using the Dot2Dot tool on chromosome 1, restricting the analysis to perfect repeats without any imperfections. From this analysis, we calculated the proportion of each motif size within the genome. These proportions were then used to guide the random selection of motifs during the simulation, ensuring that the frequency of each motif size in our simulated data mirrored the distribution observed in the human genome. The motifs were extended into longer repeats, ensuring a minimum length of 12 bases and at least 3 motifs for larger motif sizes. The repeats were interspersed with non-repetitive buffer sequences, selected from the 700 bp gaps between reported repeats in the Dot2Dot output for chromosome 1. These buffer sequences were carefully screened using both Ribbit and Dot2Dot to ensure they contained no repetitive regions. The buffer sizes and units were randomly inserted between repeats to create a more realistic genomic structure. Once the perfect repeat structures were established, we introduced imperfections in the form of substitutions, insertions, and deletions, while maintaining the overall repeat purity between 0.85 to 1. The simulated data was generated as three separate datasets, each with varying levels of impurities and a sample size ranging from 10,000 to 1 million locations. First with only perfect repeats, without any imperfections maintaining an ideal motif to motif match. Second dataset with a minimum purity of 0.90 by introducing substitutions as only form of imperfection.A substitution involves replacing one nucleotide in the sequence with another, often occurring when two motifs are highly similar but not identical. Third dataset was created through adding extra nucleotides (insertion) or removing nucleotides (deletion) from the sequence, typically occurring when there is a gap or distance between matching motifs. It had a mix of substitutions, insertions, and deletions (indels) while maintaining an overall repeat purity of at least 0.85. By introducing these mutations with a measured proportions of 80% Substitutions, 10% Deletions and 10% Insertions, we ensured that the overall sequence purity remained within the desired range while also maintaining a motif-to-motif match of at least 85%. The overall dataset was distributed into multiple chromosomes in case a sequence length was exceeding 200 Mb. Finally the script generated a fasta file while keeping a track of locations of repeats and information about the imperfections in the repeats that was written in a BED file.

## Results and Discussion

### Performance comparison based on simulated datasets

We compared Ribbit’s performance with two tools, TRF (Benson, 1999) and Dot2dot (Genovese et al., 2019). TRF, identifies repeat locations using a probabilistic model considering each periodic nucleotide match/mismatch as a Bernoulli trial. Regions with significant continuous successes or matches are tagged as TRs with calculated periodicity. Dot2dot identifies repeat regions by building a storage optimized dot-plot of the sequence, which has been used for visual identification of tandem repeats, and selects subsequences with continuous matches along the diagonals. The performance of Dot2dot is shown comparable to TRF with improvements in runtime. To compare the tool’s accuracy and runtime efficiency we used simulated repeat datasets. We created 3 sets of simulated data, the first set with only perfect repeats, the second set with substitutions as impurity with minimum purity 0.90 and the third set with substitutions and indels with minimum purity 0.85. Each dataset carried 5 samples with different numbers of locations from 10000 to 1 million. We ran Ribbit with default parameters (minimum repeat length of 12), Dot2Dot is configured with HEAVY filter mode and allowed 2 gaps, and TRF is used with the recommended settings (match=2, mismatch=5, indel=7, pm=80, pi=10, minscore=50, maxperiod=100).

Ribbit is observed to be consistently faster than Dot2Dot and TRF across all datasets. On average, the runtimes of Dot2dot and TRF were 2.5 fold and 1.4 fold longer in comparison to Ribbit. In terms of memory requirements, TRF with recommended settings is more efficient, requiring on average only 25% of the RAM used by Ribbit. Ribbit’s memory usage is comparatively higher than TRF. Meanwhile, Dot2Dot and Ribbit had comparable memory demands, with Ribbit requiring slightly higher RAM but is not observed to be consistent (Fig.1).

### Accuracy comparison

We aimed to evaluate the tools using datasets with varying levels of impurities to identify their optimal use cases and explore any edge cases. The first set of simulated datasets consisted only of perfect repeats. All tools demonstrated nearly 100% accuracy in identifying the repeat locations. The accuracy for the tools to identify the location with correct motif size was also close to 100% for Ribbit and TRF, while Dot2Dot showed approximately 93% accuracy in motif size identification. We introduced substitutions as impurities into the simulated data while maintaining an overall purity of 0.90. Ribbit, TRF, and Dot2Dot demonstrated near-perfect accuracy in identifying the locations. TRF identified 98% of the locations with the correct motif size, while Ribbit achieved 96% accuracy when allowing for 1 bp adjustment in motif size to capture a purer repeat form and Dot2Dot’s accuracy dropped significantly to 28% on this dataset. The final set of simulated data, which also included indels with a minimum purity of 0.85, yielded consistent accuracy results across all tools. Ribbit and TRF could identify motif sizes with comparable accuracy but Dot2Dot struggled to identify the correct motif size for more than 80% of the locations in the data set.

### Human data

We evaluated the performance of Ribbit and TRF on two reference human genomes: HG38 and T2T. TRF with default parameters takes days to process the whole HG38 version of the human genome. As recommended by the creators of TRF to set “-l” value to 120 which is a parameter that is to increase the expected maximum TR size runs TRF with higher RAM usage while reducing the runtime to a comparable scale. TRF is tested with two configurations: the default parameters (match=2, mismatch=5, indel=7, pm=80, pi=10, minscore=50, maxperiod=25, -l 120), and a custom parameter set (match=2, mismatch=3, indel=3, pm=80, pi=10, minscore=30, maxperiod=25, -l 120), designed to align Ribbit and TRF more closely in terms of parameter settings. For the T2T genome, the two TRF configurations showed significant differences in execution time. The default settings (2 5 7 80 10 50 100) took over 8 hours and 45 minutes, while the lenient configuration (2 3 3 80 10 30 100) required 24 hours. In contrast, Ribbit completed the run in just 4 hours and 35 minutes. TRF’s memory usage also varied between configurations, with higher memory demand linked to the “-l” parameter. For the HG38 reference, the default TRF configuration finished in 1 hour and 53 minutes, while the lenient setting took 5 hours and 42 minutes. Ribbit completed the run in 3 hours and 57 minutes. Memory usage patterns remained consistent across both genome references for both tools (Table.1).

**Table.1.** Runtime and Memory comparison of Ribbit and TRF on HG38 and CHM13-T2T Human Genome.

| Tools | HG38 |  | CHM13-T2T |  |
| --- | --- | --- | --- | --- |
|  | Runtime (minutes) | Memory usage (Gb) | Runtime (minutes) | Memory usage (Gb) |
| Ribbit | 237 | 21.4 | 275 | 21.5 |
| (Default) | 113 | 6.3 | 525 | 19.5 |
| (Custom) | 342 | 20.8 | 1,448 | 62.1 |

Ribbit provides insights into the overall TR profile of a genome. By analysing its output for the T2T and HG38 reference genomes, we could observe key differences in TR density across chromosomes [Fig.3A]. Chromosome 9 and Y show higher TR densities in the T2T reference. This is consistent with previous findings that attribute the increase in the resolution of centromeric and other complex regions [Xia, Y., Li, D., Chen, T., et al. (2024)]. We observed a peak in pentamer and hexamer density within T2T along with an increase in repeat length in these regions. [Fig.3C,D]

**Figure 3.**
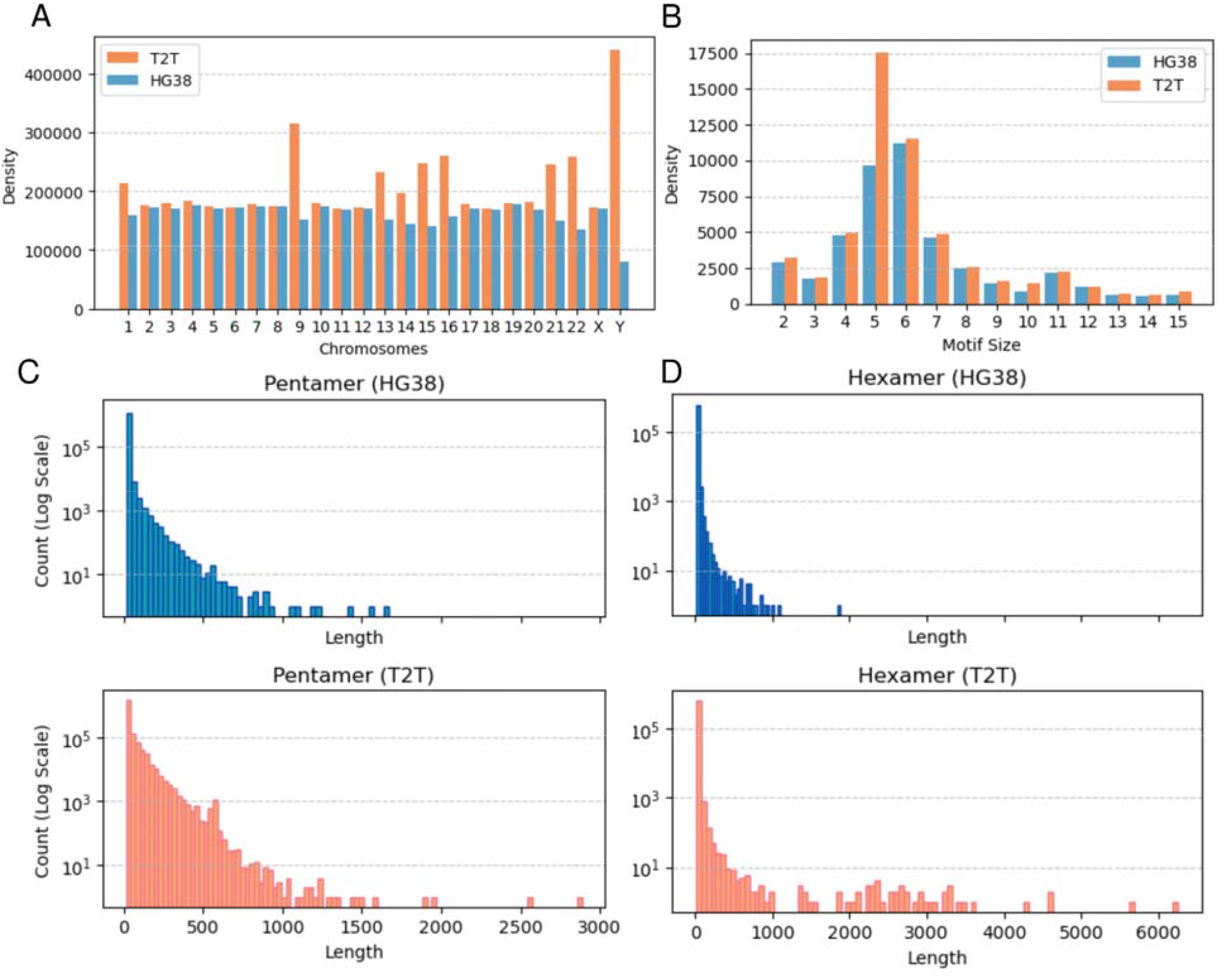
Comparative analysis for T2T and HG38 human genome references. **A)** TR density distribution across chromosomes, showing higher densities in chromosomes 9 and Y for the T2T reference genome. **B)** Density distribution of individual motifs across T2T and HG38 reference genomes. **C,D)** Length distribution histogram for pentamers revealing an increase in the length for pentamers and hexamers in T2T reference genomes.

### Representation of locations

Recent studies on the STR catalog highlight regions with diverse tandem repeats (TRs) within highly polymorphic areas known as *variation clusters* [Ben Weisburd 2024]. The authors identified 273,112 VCs that cumulatively overlapped 744,458 TRs, from a catalog of 4,863,043 TRs. These regions are best analyzed through detailed sequence analysis, yet many tools struggle to accurately resolve the individual motifs present. TRF typically annotates these regions with a single consensus motif, which limits its ability to genotype the constituent TRs separately. In contrast, Ribbit’s motif decomposition algorithm successfully identified 99.6% of the variation clusters with overlapping TRs that were more than 12 bp in length and provided a clearer resolution. These regions were represented as isolated TR regions in the BED format output and were represented as “REGION” in Ribbit’s GFF format output file. Ribbit could successfully annotate complex TR regions into a “REGION” with the relationships of overlapping or nested TRs (RSTR) detailed in the 9th column of the GFF file. TRF (2 5 7 80 10 50 100) could identify 53.7% of these variation clusters but could not resolve the regions. With TRF (2 3 3 80 10 30 100) output 91% of these clusters were identified but were represented as a consensus motif. Our comparison between TRF and Ribbit’s performance on cataloged variation clusters (Fig.3) revealed that Ribbit could effectively resolve multiple TRs where TRF had grouped them into a single motif consensus. TRF with sensitive parameters might miss out on shorter polymorphic locations. The study that mentions Variation Clusters also talks about ambiguity in defining TR start and end point, which can lead to genotype misinterpretation. For a more accurate sequence-level analysis, the authors emphasize the need to characterize TR regions in their full complexity rather than assigning a simple consensus motif as seen in TRF output [Fig 4], to better capture the inherent variability in these sequences.

**Figure 4.**
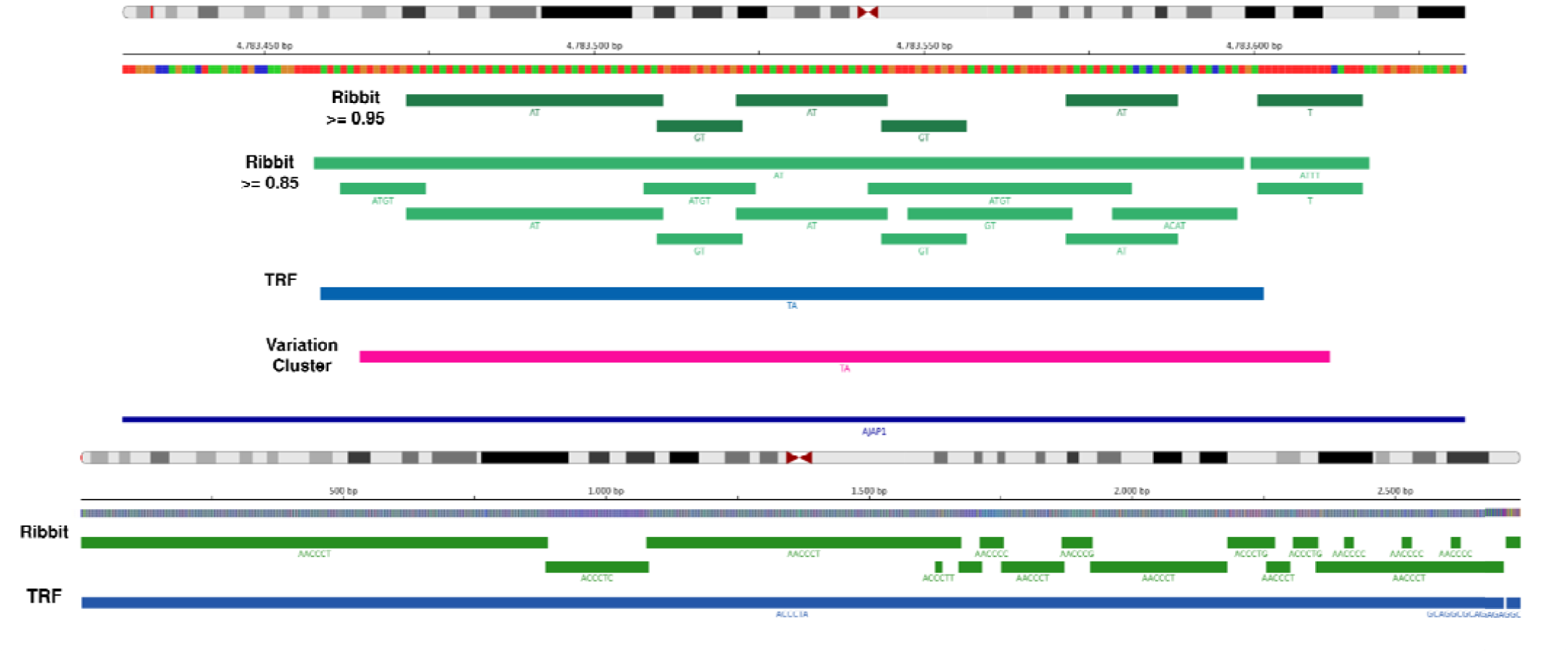
Comparative Motif Resolution for Chromosome 1 Variation Cluster and Telomere region. The regions were visualized using IGV, with distinct color bands representing different annotations: Green for filtered Ribbit calls (purity > 0.95), Light Green bands for Ribbit’s default output. Blue for TRF (parameters: 2 3 3 80 10 30 100), and Pink highlighting the variation cluster regions **A)** chr1:4783464-4783611, featuring overlapping AT and GT repeats alongside a T monomer. **B)** Telomeric region in Chromosome 1.

We also looked at telomeric regions in the T2T reference for different chromosomes. (Fig.4), that showed presence of nested repeats inside AACCCT repeat region and short stretches of other motifs overlapping the telomeric repeat. The minimal repeating unit of a telomere varies from species to species and such analysis can help us to explore the evolutionary divergence of telomeres. Ribbit performs a pairwise alignment while accounting for interruptions that introduce slight mismatches, prioritizing the motif that is repeated most frequently. When a clear representative motif for a significant part of a region cannot be found, Ribbit iterates its algorithm to identify the most suitable motif for that region. This iterative approach allows Ribbit to resolve complex TR regions while maintaining relationships between repeats and reporting interruptions. Many of these regions are also part of Variation Clusters in the TR catalog. Another benefit of using pairwise alignment is that it provides a global picture of all the variations (substitutions, indels) across the entire repeat region. It’s easier to understand the distribution of imperfections throughout the sequence. Motif to motif alignment could fragment the repeat while analyzing and might provide an disjointed picture of the repeats as observed in Dot2Dot outputs. In case of larger variations, such as deletion or insertions that might affect multiple motifs, are easier to capture with pairwise alignment of repeat regions with their corresponding pseudo perfect repeat sequence. Pseudo perfect repeat acts as a standard sequence for comparison and gives a perspective to understand how the repeat region has deviated from its “ideal” form. The calculations for the degree of variation in a TR becomes easier.

### Ribbit’s output

Ribbit not only outputs individual repeat locations in a BED file but also defines the relationships between repeats by providing a GFF file. It generates a well-structured, informative output, including a CIGAR string that details TR characteristics, such as the location and type of imperfections. This CIGAR string offers valuable insights into the nature of imperfections within the tandem repeats, facilitating detailed analysis of TR patterns and variations. Ribbit’s GFF output format further enhances the examination of nested or compound TR relationships, enabling users to fully explore and understand the complex dynamics of these repeat regions. Ribbit is built to perform repeat identification on reference genomes (fasta formats), that fairly represents the repeat landscape of any genome.

## Conclusion

As we developed our tool for identifying tandem repeats (TRs) in the genome, we explored how the definition of a TR has evolved over time and identified key aspects of TR regions worth examining. Early tools defined a TR as any motif that repeats more than twice in succession. Later, this definition expanded to include regions with periodic repetition and minor motif variations. Now, for genotyping purposes, a more detailed understanding of these regions is essential. Many previous approaches to capture imperfections in TRs failed to account for all types of variation. The discovery of novel TR regions in the human genome continues because of improved algorithms and complete genome assemblies like CHM13-T2T. This presents an exciting opportunity to study TR regions more comprehensively.

We compared the performance of Ribbit and TRF, noting that Ribbit performs much the same as TRF with custom configurations but with greater efficiency and less redundancy. In our analysis of TR density per 1 Kbp across the human genome, both tools produced closely aligned results. However, Ribbit represents the repeat motifs more precisely and resolves the overlapping or nested repeats. TRF with its default settings tends to ignore valid TR locations while with the custom settings more than 35 % of the genome is annotated as repetitive. Ribbit’s default settings, on the other hand, offer a more detailed and less redundant TR profile. Ribbit allows users to explore the repetitive nature of any DNA sequence with its customizable parameters and easier operation.

## Code availability

- https://github.com/SowpatiLab/ribbit Ribbit repository.
- https://github.com/SowpatiLab/ribbit/tree/main/data_simulation Script used to generate simulated datasets.
- https://github.com/SowpatiLab/ribbit/tree/main/scripts Script to convert BED to GFF format.

## Notes

### Competing Interest Statement

The authors have declared no competing interest.

