## Supplementary for "Ribbit: Accurate identification and annotation of complex tandem repeat sequences in genomes"

### Supplementary Information
### Supplementary Methods

##### Generating Shift XOR bitsets

The core of Ribbit's algorithm is the generation of the shift XOR bitsets. To compare every base with its periodic neighbor we do an XOR between the original sequence bitset and the sequence shifted by $u$ bits. X OR yields a bitset with 1s denoting mismatches and 0s denoting matches. The bits are flipped using a NOT operation for 1s to denote matches and 0s denote mismatches. The resultant bitset is referred to as the shift XOR bitset and shortly as *SXOR*. This is done individually for both the left and right bitsets which are then combined using AND operation. Based on the input of the desired motif sizes, ribbit generates the *SXOR* for all motif sizes.

##### Sliding window approach

The shift XOR bitsets generated for various motif sizes undergo further analysis to pinpoint potential repeated units, referred to as 'seeds'. This identification process involves employing a heuristic approach. Initially, a sliding window of length 8 bits is utilized, moving one bit at a time along the bitsets. For each window, the number of set bits (1s) is counted. If the bit count is at least 50% (equivalent to at least four 1s), the window moves forward. Additionally, the start and end indexes of each identified seed are recorded. These indexes provide crucial information regarding the position of potential tandem repeats within the sequence. Window start is updated on encountering appropriate bitcount and the end index of the seed is updated when the next window, adjacent to the window with a lower bit count, is checked. This ensures that the end of the seed accurately reflects the span of the potential repeat region. The final start and end index is updated as the seed bitset position.

##### SIMD implemented Smith Waterman alignment

To gain the insights into the impure form of TRs, the algorithm compares it with the idealized counterpart. In this process, a pseudo-perfect TR, with a length equal to the seed sequence, is generated by repeating the most frequent motif identified earlier. This pseudo-perfect TR serves as a reference for evaluating the impurities within the original seed sequence. Ribbit employs a Single Instruction Multiple Data (SIMD) implementation of the Smith-Waterman alignment, optimizing the alignment process for thousands of seeds within seconds. SIMD, a computer architecture feature available on x86 CPUs that facilitates parallelization of the algorithm at the instruction level enabling simultaneous execution of similar operations on multiple datasets. The integration of SIMD significantly enhances the algorithm's efficiency and the use of SIMD Smith-Waterman (SSW) C/C++ library further accelerates the alignment process ~50 times compared to the standard SW library. The alignment reports SW score, alignment location and traceback path (CIGAR) along with the suboptimal alignment and location heuristically. [3]

##### Measures of purity

Ribbit allows for identification of STRs with three different purity measures which are implemented independently. The first measure is fraction purity which is the number of matching nucleotides of the STR sequence when aligned with a perfect repeat sequence of STR.

##### Processing CIGAR

Ribbit processes CIGAR string that was generated after the alignment of seed sequence with pseudo perfect repeat sequence to understand the characteristics of that TR. CIGAR is split into components as lengths and types such as Soft Clips (‘S’), Mismatch (‘X’), Insertion (‘I’), Deletion (‘D’) and Match (‘M/=’). The first measure is fraction purity which is the number of matching nucleotides of the STR sequence when aligned with a perfect repeat sequence of STR. In pruning, a compressed cigar is produced after looking for the continuous mismatches, trimming soft clips and adjusting repeat start and end indices.

#### Supplementary Figures

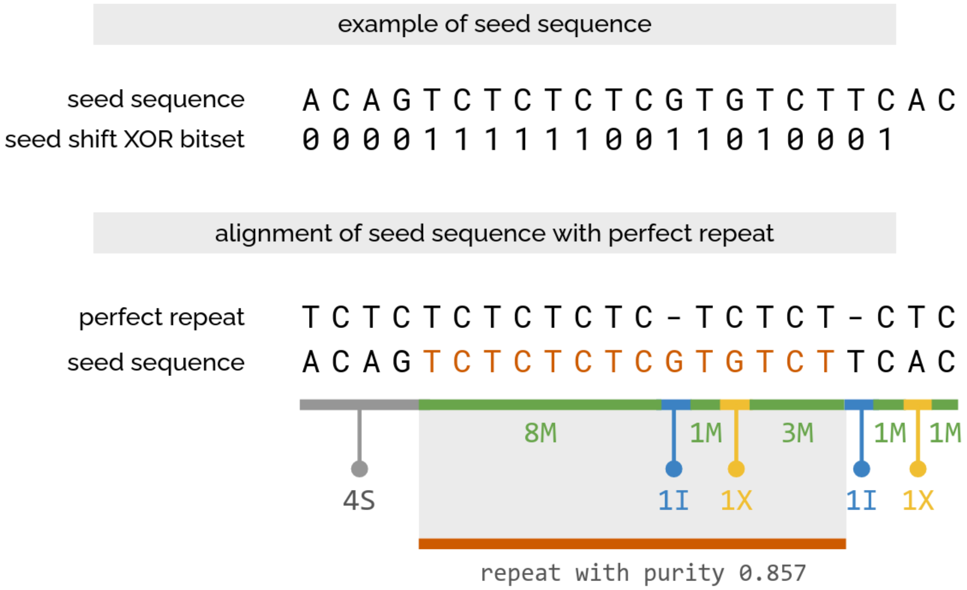


**Supplementary Figure 1.** Alignment of the seed sequenct post selection with a pseudo perfect repeat. The impurities in a repeat sequence are identified at this step.

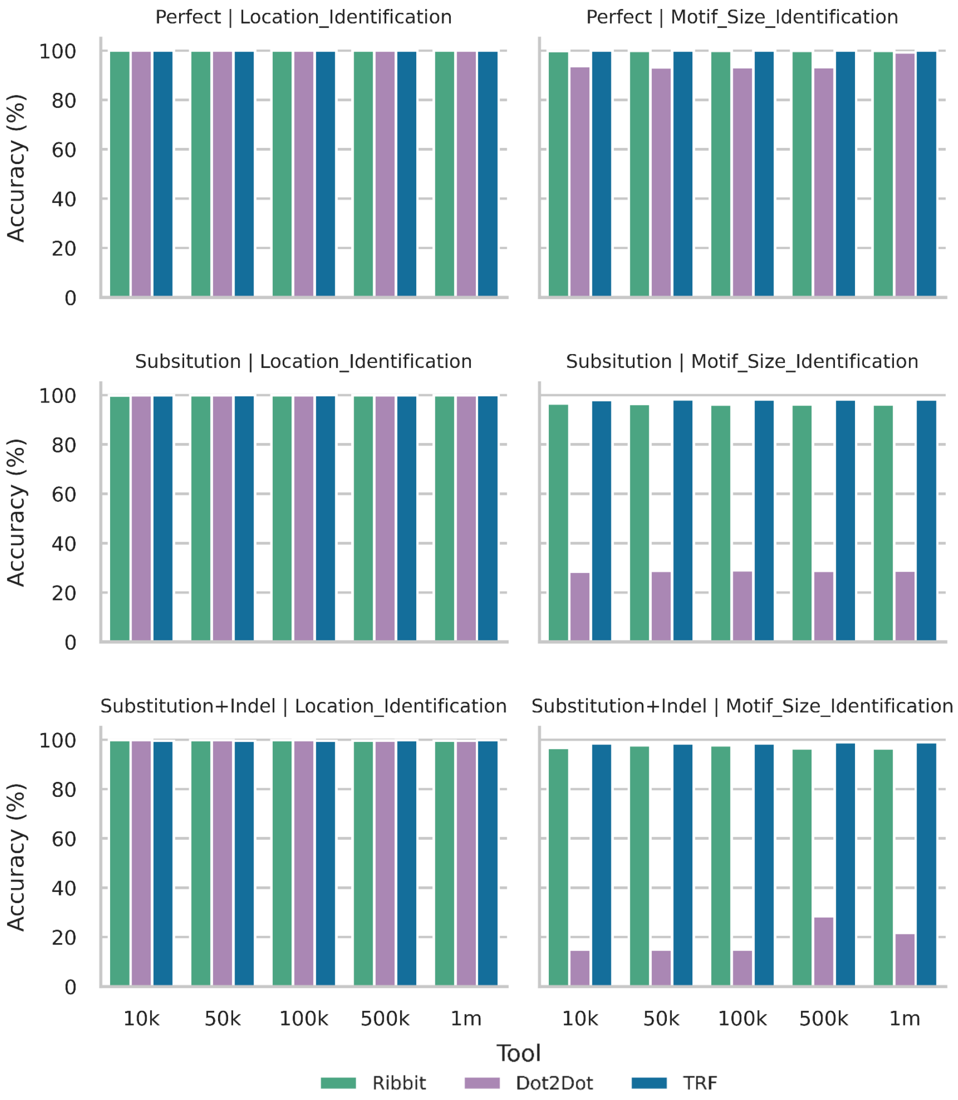


**Supplementary Figure 2,** Accuracy comparison between Ribbit, Dot2Dot and TRF across all 3 simulated datasets for all (10k-1m) sample sizes.

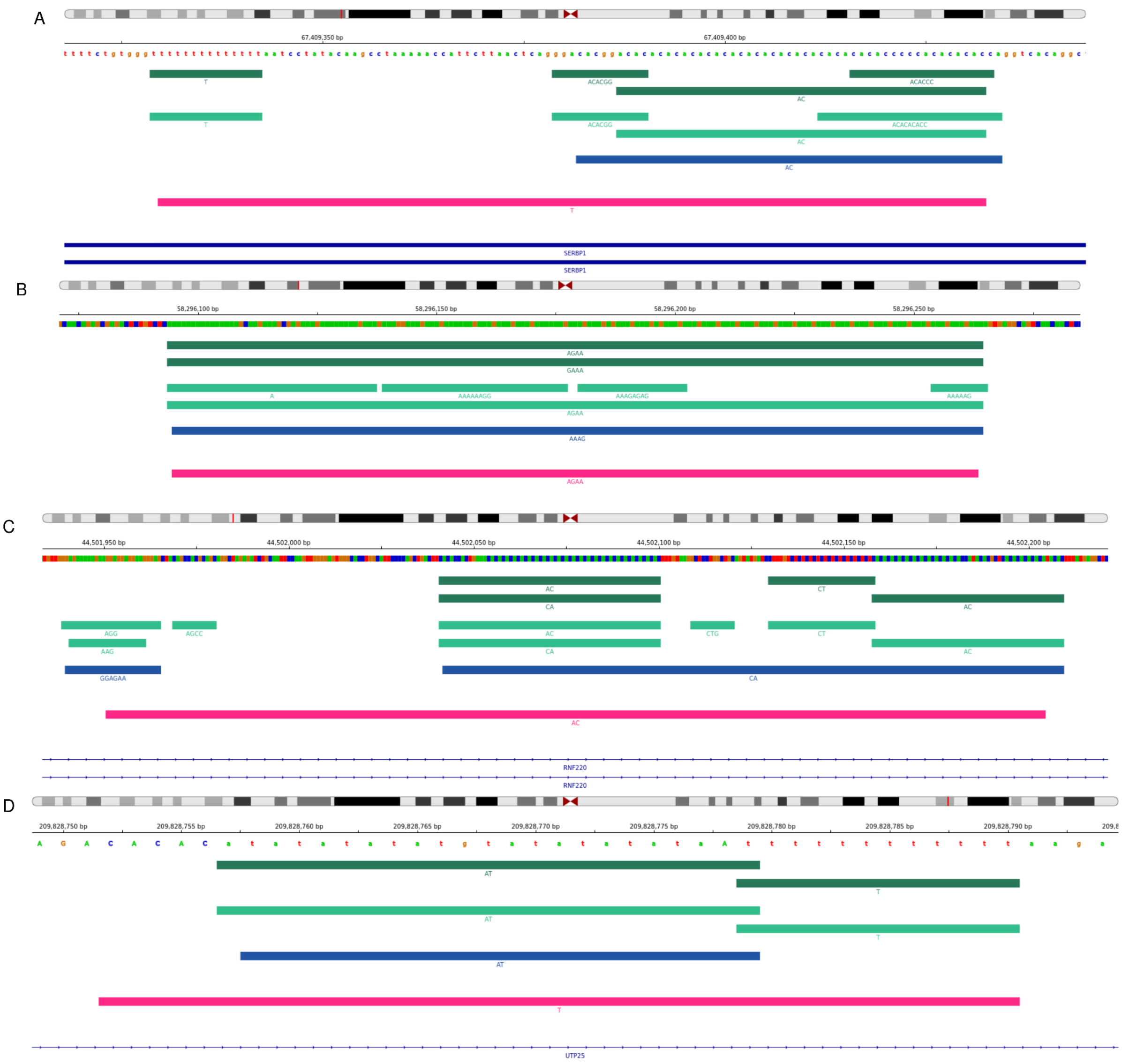


**Supplementary Figure 3.** The regions were visualized using IGV, with distinct color bands representing different annotations: Green for filtered Ribbit calls (purity > 0.95), Light Green bands for Ribbit's default output. Blue for TRF (parameters: 2 3 3 80 10 30 100), and Pink highlighting the variation cluster regions A) chr1:67,409,329-67,409,432. B) chr1:58,296,094-58,296,263 C) chr1:44,501,950-44,502,204 D) chr1:209,828,751-209,828,790
